# Long divergent haplotypes introgressed from wild sheep are associated with distinct morphological and adaptive characteristics in domestic sheep

**DOI:** 10.1101/2022.05.17.492311

**Authors:** Hong Cheng, Zhuangbiao Zhang, Jiayue Wen, Johannes A. Lenstra, Rasmus Heller, Yudong Cai, Yingwei Guo, Ming Li, Ran Li, Wenrong Li, Sangang He, Jintao Wang, Junjie Shao, Yuxuan Song, Lei Zhang, Masum Billah, Xihong Wang, Mingjun Liu, Yu Jiang

**Affiliations:** Key Laboratory of Animal Genetics, Breeding and Reproduction of Shaanxi Province, College of Animal Science and Technology, Northwest A&F University, Yangling 712100, China; Faculty of Veterinary Medicine, Utrecht University, Utrecht, The Netherlands; Shenzhen Branch, Guangdong Laboratory of Lingnan Modern Agriculture, Genome Analysis Laboratory of the Ministry of Agriculture and Rural Affairs, Agricultural Genomics Institute at Shenzhen, Chinese Academy of Agricultural Sciences, Shenzhen, China; Section for Computational and RNA Biology, Department of Biology, University of Copenhagen, DK-2100 Copenhagen, Denmark; Key Laboratory of Ruminant Genetics, Breeding & Reproduction, Ministry of Agriculture, China; Key Laboratory of Animal Biotechnology of Xinjiang, Institute of Biotechnology, Xinjiang Academy of Animal Science, Urumqi, Xinjiang 830026, China

**Keywords:** domestic sheep, introgression, horn status, ear morphology

## Abstract

The worldwide sheep population comprises more than 1000 breeds. Together, these exhibit a considerable morphological diversity, which has not been extensively investigated at the molecular level. Here, we analyze whole-genome sequencing individuals of 1,098 domestic sheep from 154 breeds, and 69 wild sheep from seven *Ovis* species. On average, we detected 6.8%, 1.0% and 0.2% introgressed sequence in domestic sheep originating from Iranian mouflon, urial and argali, respectively, with rare introgressions from other wild species. Interestingly, several introgressed haplotypes contributed to the morphological differentiations across sheep breeds, such as a *RXFP2* haplotype from Iranian mouflon conferring the spiral horn trait, a *MSRB3* haplotype from argali strongly associated with ear morphology, and a *VPS13B* haplotype probably originating from urial and mouflon possibly associated with facial traits. Our results reveal that introgression events from wild *Ovis* species contributed to the high rate of morphological differentiation in sheep breeds, but also to individual variation within breeds. We propose that long divergent haplotypes are a ubiquitous source of phenotypic variation that allows adaptation to a variable environment, and that these remain intact in the receiving population due to reduced recombination.

## Introduction

Domestic sheep (*Ovis aries*) descends from Asiatic mouflon(Ryder and Mason 1981; Zeder 2008) approximately 11,000 years ago in southeastern Anatolia of Turkey. As many as 1,400 different breeds (Scherf 2000) exhibit a remarkable phenotypic diversity in response to selection pressures from a diverse range of environments as well as to human selection. The wild sheep from the *Ovis* genus (snow sheep, *O. nivicola*; bighorn, *O. canadensis*; thinhorn, *O. dalli*; argali, *O. ammon*; urial, *O. vignei*; Asiatic mouflon, *O. orientalis* and European mouflon, *O. musimon*) are widely distributed along the East-West axis from Eurasia to North America(Rezaei, et al. 2010), with evolutionary relationships matching their biogeographic history(Rezaei, et al. 2010; Cao, et al. 2020; Lv, et al. 2021).

Recent studies have documented introgressions from various wild relatives into domestic sheep (Barbato, et al. 2017; Hu, et al. 2019; Cao, et al. 2020; Li, Yang, Shen, et al. 2020; Lv, et al. 2021). However, most reports focused on isolated cases of gene flow between two sympatric *Ovis* species, e.g. the introgression from European mouflon into European domestic breeds(Barbato, et al. 2017; Cao, et al. 2020), from Iranian mouflon into domestic sheep(Li, Yang, Li, et al. 2020), from Asiatic mouflon to Grey Shirazi, and from argali to Tibetan sheep(Hu, et al. 2019). Moreover, some of these studies based on the sheep 50K SNP BeadChip considered only a limited number of variants to evaluate the introgression proportions(Barbato, et al. 2017; Hu, et al. 2019). Given these recurrent findings of interspecies introgression, it would be preferable to jointly infer the magnitude of such introgression across the whole genus, as pairwise introgression results can be biased by ignoring the presence of other introgression events in such reticulated evolution scenarios. Nonetheless, these studies have yielded interesting evidence for introgression of functional genes, such as the *HBB* locus as adaptation to the high-altitude of the Qinghai-Tibetan plateau.

Here we used 1,167 whole-genome resequenced sheep (**Fig.1, supplementary Table S1** and **S2**) with 156 samples were newly generated (**supplementary Table S2**). We phased the genomes into haplotypes for an integrative analysis of the introgression from different wild sheep species. We further collected genotypes and phenotypes from East-Friesian sheep × Hu Sheep F2 hybrids to annotate the potential functional impact of various introgression signals. Our results provide further insight into the reticulated history of sheep evolution and particularly into the role of divergent haplotypes in the phenotypic diversity.

**Fig. 1.**
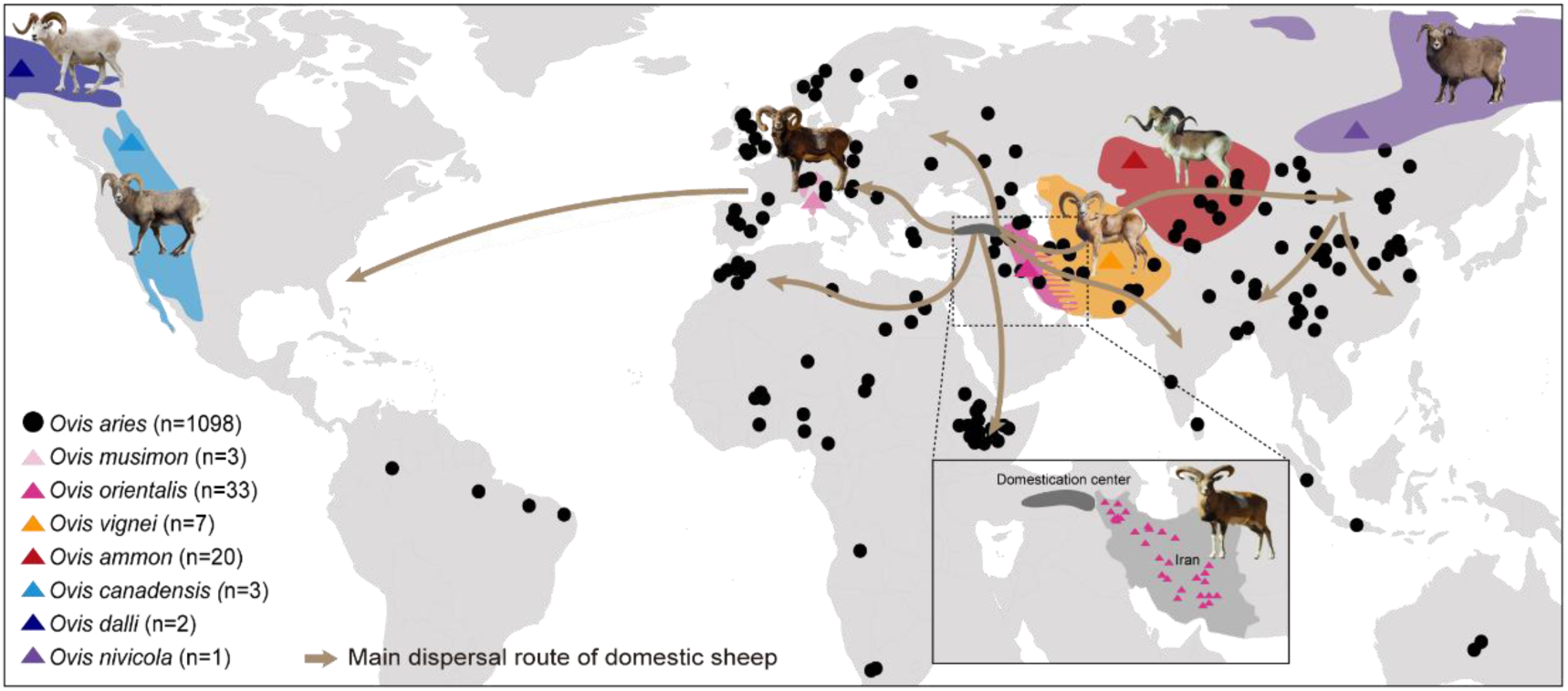
Locations of different geographically wild *Ovis* species and diverse domestic sheep populations used in this study. The colored blocks show the geographic distributions of the wild species. And each black dot represents a domestic breed. The dark grey block means the domestication center of sheep, and the solid lines represent dispersal routes of domestic sheep out of their domestication areas.

## Results

### Genetic variant data and phylogenetic relationships of *Ovis* genus

To investigate the phylogeny and population differentiation of *Ovis* species, we collected and generated a whole genome SNP dataset from 1,167 individuals comprising 1,098 domestic sheep across the geographic distribution of 154 breeds and 69 samples of their seven wild relatives (**supplementary table S1**). After aligning reads to the Oar_v4.0 (GCF_000298735.2) and quality control, a total of 83,386,953 SNPs were detected.

A whole-genome maximum likelihood (ML) phylogenetic tree revealed that European mouflon is intermediate between Asiatic mouflon and domestic sheep (**Fig. 2A, supplementary Fig. S1**). This is in agreement with their descent from the ancestral population of European domestic sheep, which were then subsequently replaced by the first domestic wool sheep populations. Domestic sheep was much closer to the Iranian mouflon located in western Iran (**supplementary Fig. S2**), near to the domestic center. The evolutionary relationships among other wild sheep were consistent with the topology inferred by mtDNA sequences(Rezaei, et al. 2010). Principal component analysis (PCA) further divided *Ovis* species intro three separate clusters (1) *O. nivicola, O. canadensis* and *O. dalli*; (2) *O. ammon*; (3) *O. vignei, O. orientalis, O. musimon* and *O. aries* (**Fig. 2B, supplementary Fig. S3**). The PCA of mouflon and domestic sheep as well as the ADMIXTURE pattern at k⩾7 (**Fig. 2F, supplementary Fig. S4**) show a differentiation of eastern and western Iranian mouflon according to their geographic origin (**Fig. 2B**). Moreover, the PCA confirms the relatively close relationship of western Iranian mouflons and domestic sheep. Both PCA and ADMIXTURE at k=8 reveal a correlation of genetic clustering and geographic distances for domestic sheep (**Fig. 2D** and **2F, supplementary Fig. S3** and **S4**). Samples from China were subdivided into three groups (**Fig. 2E** and **2F**), CN_YNS (Yunnan sheep), CN_TIB (Oula, Prairie Tibetan, Valley Tibetan) and CN_NOR (Small tailed Han sheep, Cele black sheep, Hu sheep, Tan sheep, Bayinbuluke sheep and Ujimqin Sheep).

**Fig. 2.**
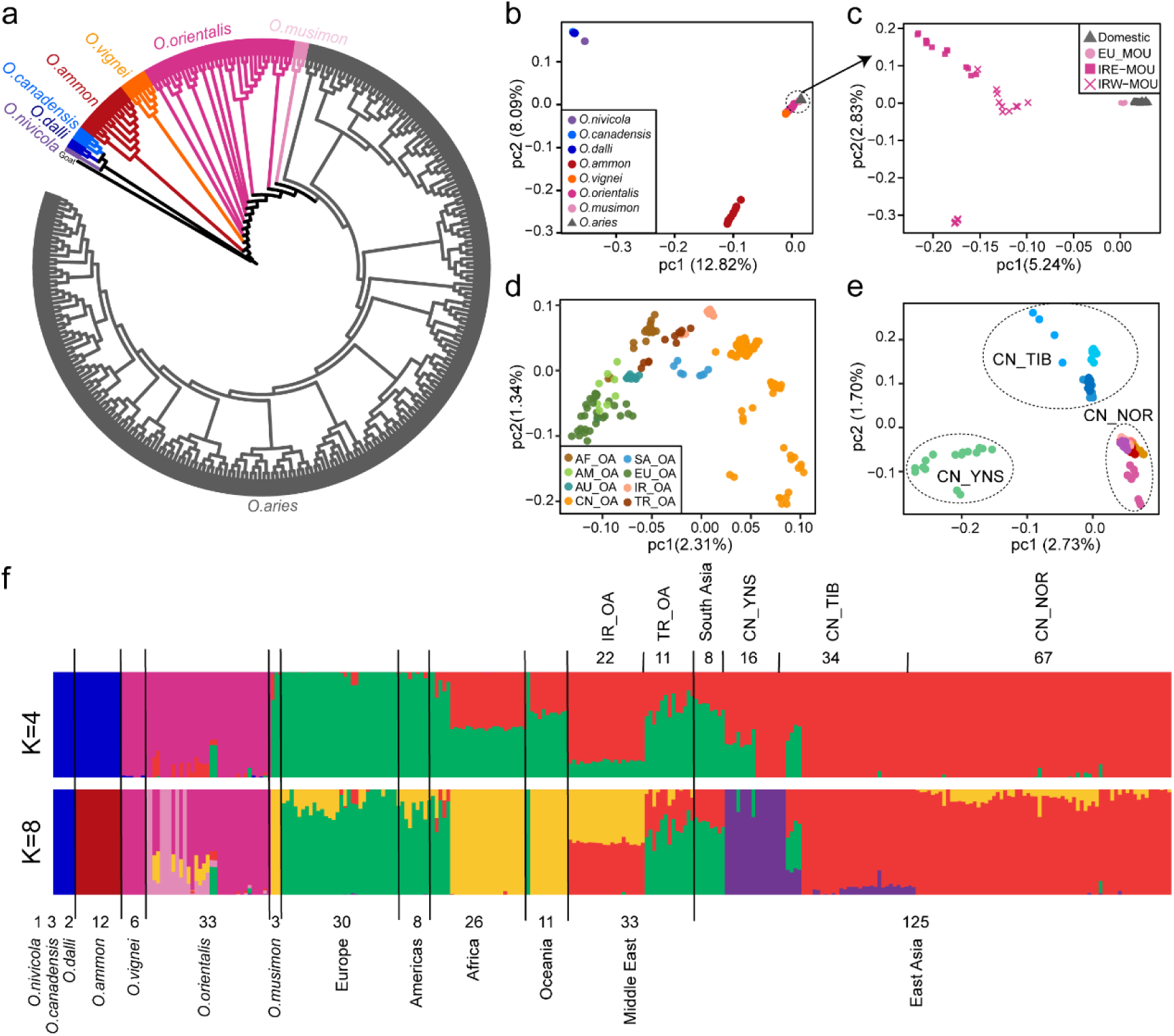
Phylogenetic analysis and population genetic structure. (A) A maximum likelihood (ML) phylogenetic tree of 293 representative samples covering all species of *Ovis* genus with Goat (GCA_000317765.2) as an outgroup. The tree was built with 100 bootstraps using a total of 332,990 4DV sites. (B-E) PCA analysis of wild and domestic sheep (B), Iranian and European mouflon and domestic sheep (C), domestic (D) and Chinese sheep (E), respectively. EU_MOU, European mouflon; IR-MOU, mouflon sheep from Iran. AF_OA, AM_OA, AU_OA, CN_OA, SA_OA, EU_OA, IR_OA, TR_OA, separately represent domestic sheep from Africa, America, Oceania, China, south Asia, Europe, Iran and Turkey. CN_TIB, CN_YNS, CN_NOR for domestic sheep from Tibet, Yunnan, Northern China. (F) ADMIXTURE results for k=4 and k=8.

### Introgressions from wild relatives into domestic sheep

To evaluate the admixture proportion and locate the putative introgressive fragments in domestic populations from their wild relatives, we performed local ancestry inference (LAI) method program LOTER for each fully phased sheep genome. The bighorn, thinhorn, argali, urial, Iranian mouflon and European domestic sheep were used as source populations. European domestic sheep, which has not been in contact with the Asian wild sheep populations following their divergence, shared only few alleles with wild species (**supplementary Fig. S5b**) and was used as the non-introgressed reference population. The European mouflon was not tested as a source population due to its close relationship with domestic sheep (**Fig. 2**), which would confound the detection of introgression from the other wild sheep species.

In order to distinguish the putative signals of introgression from shared ancestral polymorphisms (incomplete lineage sorting, ILS), we calculated the expected length L of ILS tracts (see Materials and Methods) and removed the inferred introgressed segments with a length < L. This could remove some short introgressed regions, but is justified by the expectation that introgressed regions are considerably longer as they had less time to be broken up by recombination. In addition, we were mostly concerned about long introgressed haplotypes in the present study.

Using the filtered results, we calculated the genome-wide proportions of admixture. We detected an average of 10,036 segments (5,600-13,057), in total corresponding to an average of 180 Mb of wild *Ovis* sequence (range 96-224 Mb, SD=23Mb) for each haploid domestic sheep genome. The average proportions of domestic sheep genome from Iranian mouflon, urial, argali, bighorn and thinhorn sheep were 6.8% (3.8-8.5%), 1.0% (0.5-1.4%), 0.2% (0.07-0.3%), 0.03% (0.01-0.05%) and 0.01% (0.006-0.022%), respectively (**Fig. 3A, supplementary Fig. S6-7**), values that are similar to those previously reported for sympatric wild-to-domestic introgression.5 The introgressed proportions varied considerably across wild donor species, in particular between Iranian mouflon and the other wild species (**Fig. 3A**). East Asian domestic sheep has a relatively strong introgresssion from urial and argali (**Fig. 3A**), consistent with their biogeographic history.

**Fig. 3.**
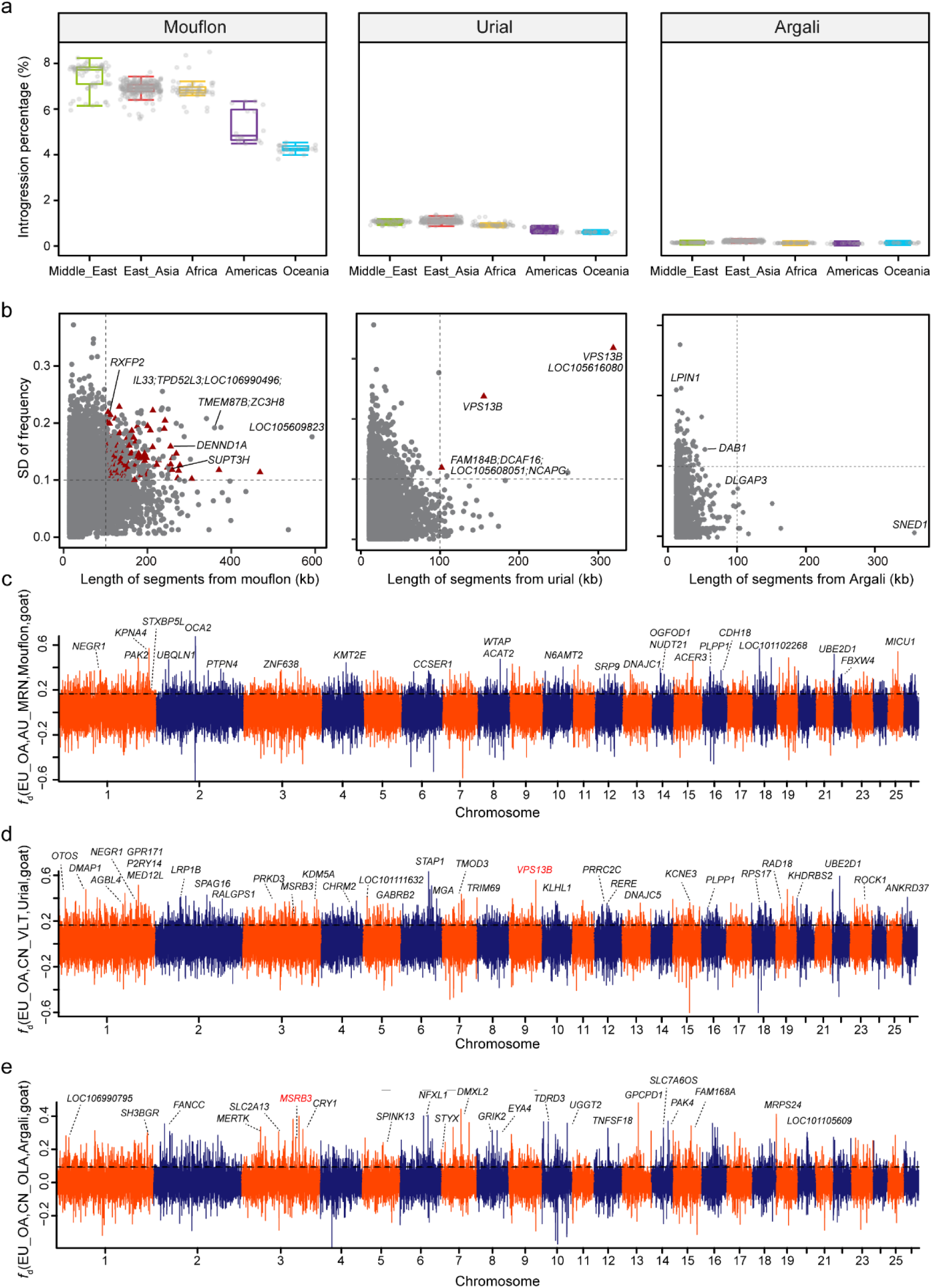
Genome-wide introgression evaluation and potential adaptive introgression. (A) The proportions of introgressed sequences from wild relatives (Iranian mouflon, urial and argali) identified in each domestic population. Each dot indicates a phased haploid. (B)Joint distribution of length for introgressed tracks (x axis), SD for introgressed haplotype frequency among distinct populations (y axis). Red triangles indicate tracts with top 1% *F*_ST_ values between at least one of the 16 domestic meta-populations and Iranian mouflon. (C-E) Manhattan plot of values showing the introgression signals from Iranian mouflon to Australian Merino (C), from Urial to Valley Tibetan sheep (D) and from Argali to Tibetan Oula sheep (E). The horizontal dashed line indicated the *P*<0.05 cutoff.

We further computed the modified f-statistic value(Martin, et al. 2015) for each 50-kb window with a 20-kb step across the genomes in the form *f*_*d*_ (European domestic sheep, domestic population; wild source of introgression, goat) (**Fig. 3C-E**). We grouped the domestic samples into 16 focal populations (see Materials and Methods). For each population, the regions with significant *f*_*d*_ values (*P* < 0.05) were defined as potentially introgressed regions(Teng, et al. 2017; Hu, et al. 2019). We further estimated *d*_XY_, phylogenetic trees and haplotype networks to corroborate the signals of introgression in specific regions.

### Selection and adaptive signatures for introgressed segments

We focused on those introgressed haplotype blocks that are conserved within but not across populations, since they are most likely involved in population differentiation and adaptation to local habitats or selection (Janzen, et al. 2019). For this, we calculated allele frequencies of the introgressed fragments in 16 domestic meta-populations (see Materials and Methods). Next, we inferred 483 mouflon, 5 urial and no argali outlier haplotypes, putatively introgressed on the basis of their length (⩾100 kb), their total frequency (⩾ 0.05) and frequency variation in the 16 meta-populations (> 0.1 standard deviation)(**Fig. 3B**).

In order to detect fragments that are possibly involved in selection, we plotted *F*_ST_ for each of the 16 meta-populations to the Iranian mouflon across the genome in 50-kb windows with a 20-kb step size. With a top 1% *F*_ST_ cutoff and joining the windows that were separated by a distance of ⩽50 kb, we obtained 2,101 non-overlapping selective sweep regions. These blocks were slightly but significantly longer than the general blocks (**supplementary Fig. S8**). For the 488 mouflon and urial introgression outliers above, 116 and 3 were overlapped with these selective sweep regions (**Fig. 3B**) and were here studied on more detail.

As expected, introgressed haplotype blocks are unevenly distributed among the domestic populations with a clear geographic signal. For instance, in region chr2: 109,998,387-110,183,036, the frequency of the Iranian mouflon derived haplotype is high in Iran local breeds (0.60) and Tan sheep (0.61), but very low (0.00) in Australian Merino and several Chinese breeds such as Hu sheep, Ujimqin Sheep, and Valley Tibetan sheep. The longest introgressed urial haplotype chr9:77,117,407-77,437,296 has a high frequency in Tibetan sheep including Oula (0.75), Prairie (0.90) and Valley Tibetan (0.80) and is almost entirely absent in sheep from Africa, America, Oceania and the Middle East. Overall, these putatively introgressed regions contained 891 genes, of which 883 and 8 were within the haplotypes derived from Iranian mouflon and Urial, respectively. It is noteworthy that within subset of introgressed regions that we identify as being under selection, several genes have been associated with morphological traits, particularly in facial shape. *RXFP2* was strongly associated with sheep horn morphology (Johnston, et al. 2011; Kijas, et al. 2012), and *SUPT3H* was reported to be associated with nose bridge breadth(Adhikari, et al. 2015), nose morphology(Claes, et al. 2018), chin dimples(Pickrell, et al. 2016) and forehead protrusion (Bonfante, et al. 2021). *MSRB3* had been identified as a candidate gene for external ear morphology in pig, dog, goat and sheep (Boyko, et al. 2010; Webster, et al. 2015; Wei, et al. 2015; Zhang, et al. 2016; Zhang, et al. 2017; Chen, et al. 2018a; Kumar, et al. 2018; Paris, et al. 2020). Furthermore, several other genes (e.g., *STXBP5L, DENND1A, VPS13B*) were identified in GWAS studies of human facial shape analyses (Claes, et al. 2018; Bonfante, et al. 2021; White, et al. 2021).

### Introgressed *RXFP2* affects horn status

There are three main types of horn status in sheep (1) horned males and females (“horned”); (2) horned males, polled females (“sex-specific”); (3) polled males and females (“polled”) (Castle 1940; Dolling and CHS 1961). A previous study indicated that the “horned” haplotypes in Tibetan sheep within *RXFP2* was most likely introgressed from argali (Hu, et al. 2019). However, in our study the same region, not argali (chr10: 29,435112-29,481,215) was detected as introgression from Iranian mouflon (**Fig. 3B**). Furthermore, we found that this introgressed region has clear signatures of selection in breeds with different horn status (**Fig. 4A, supplementary Fig. S9**). LAI indicates that most haplotypes in breeds with horn status (3) contain haplotypes most closely related to those of Iranian mouflon (**Fig. 4B-D**), pointing to a possible origin of this phenotype from this wild sheep species. We further investigated in detail the pattern of haplotype variation in this region.

**Fig. 4.**
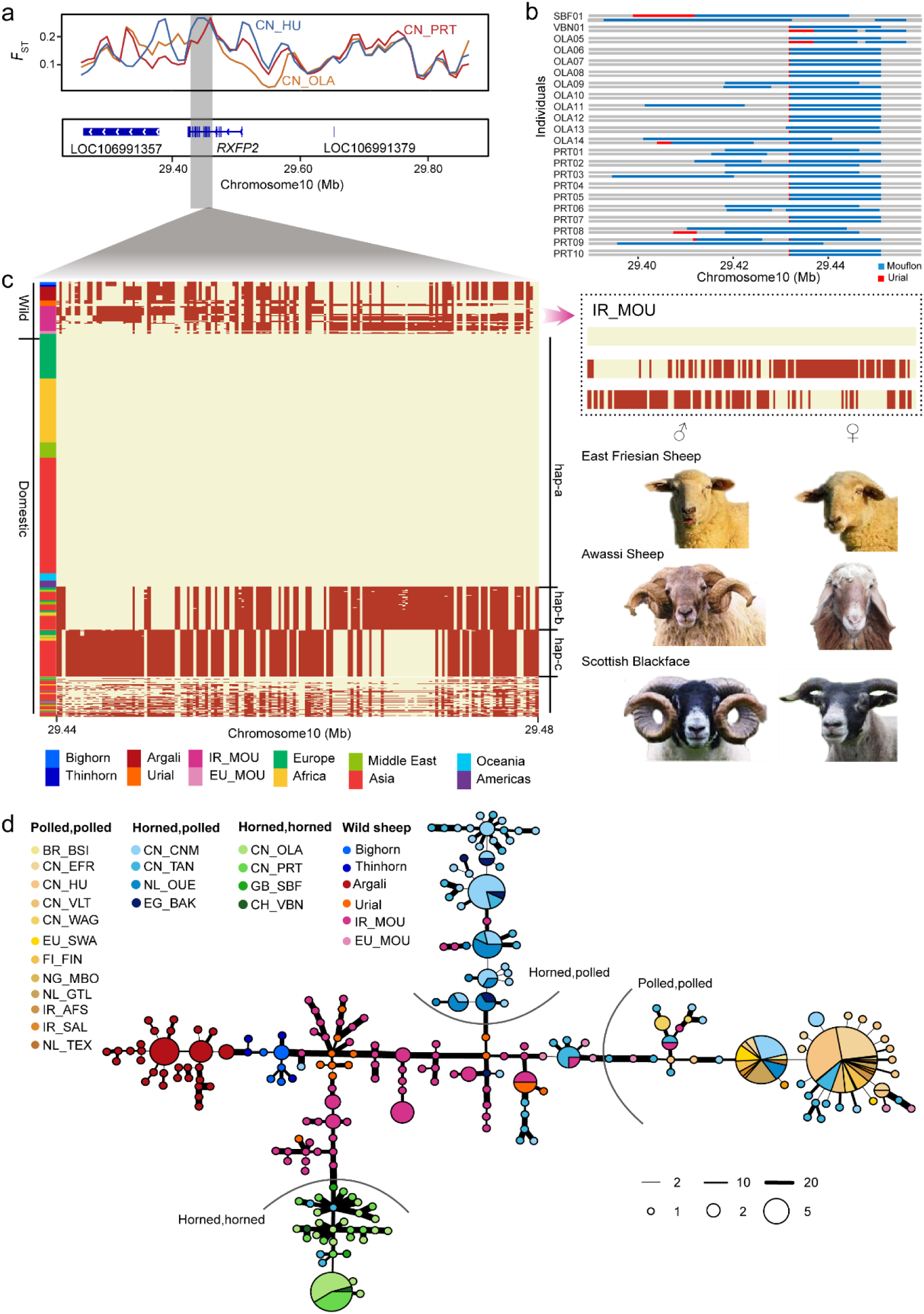
Identification and haplotype resolution of introgressed fragment at the *RXFP2* locus. (A) Distribution of the pairwise fixation index (*F*_ST_) values between Iranian mouflon and domestic sheep populations for each 50-kb window (top track). Gene annotations in the selected region in Oar_v4.0 are indicated at the bottom. (B) LAI within *RXFP2* in Valais Blacknose, Scottish Blackface, Oula and Prairie Tibetan sheep illustrating mosaic patterns of source population. (C) The haplotype pattens of *RXFP2* introgressed region (Oar chr10: 29,436,086-29,466,717). Each row is a phased haplotype, and each column is a polymorphic SNP variant. The refence and alternative alleles are indicated by light yellow and red, respectively. The haplotypes present in Iranian mouflons are indicated separately. (D) A haplotype network generated by the R software package PEGAS based on 221 SNPs of 666 haplotypes.

Haplotype patterns in this region across all 1,167 sheep showed three major highly divergent haplogroups (hap-a, hap-b and hap-c), with a few other diverse or recombinant haplotypes at low frequency (**Fig. 4C, supplementary Fig. S10**). Hap-a is the dominant haplotype in domestic sheep (**Fig. 4C, supplementary Fig. S10**) and is completely fixed (frequency=1) in Finnsheep (n=12), Gotland (n=10), Waggir (n=9), Afshari (n=6), and East Friesian sheep (n=10) (**supplementary Fig. S11, supplementary Table S4**), all of which have the “horned” phenotype. Intriguingly, hap-a is present as heterozygotes in two Iranian mouflon samples (**Fig. 4C**). ILS can virtually be ruled out because of a low probability (0.00493 for a 46,103 bp haplotype). We concluded that polledness likely occurred in wild sheep progenitors, possibly as recessive trait, and rapidly became widespread in domestic sheep because it was under strong selection in a domesticated setting.

Hap-b is generally found at high frequency in breeds with the “sex-specific” horn phenotype, including Chinese Merino (25/40, 0.625), Ouessant (12/20, 0.6) and Barki sheep (5/6, 0.83). Hap-c in contrast is usually at high frequency in breeds with the “horned” phenotype, including Oula, Prairie Tibetan, Valais Blacknose and Scottish Blackface sheep (**supplementary Fig. S11** and **supplementary Table S4**).

**Fig. 4D** shows a network of intact non-recombined haplotypes in the ∼46-kb region around *RXFP2* from wild and domestic sheep. The network suggests that haplogroups corresponding to Hap-a, Hap-b and Hap-c, respectively, are all linked to haplotypes that occur in Iranian mouflon. However, in the network and in the ML tree (**supplementary Fig. S12**), Hap-a and Hap-b haplotypes of mouflon are intermingled with those of urial, so it cannot be ruled out that the introgressed fragments originate from urial and were introgressed into domestic sheep via the mouflons. The Valais Blacknose and Scottish Blackface (European domestic breeds) haplotypes were assigned to the “horned” phenotype cluster, validating the earlier introgressed time for this locus as well. Nucleotide difference between the Iranian mouflon haplotypes and hap-c (**Fig. 2B**) suggest that hap-c was introgressed from a mouflon sub-population of Asiatic mouflon that has not yet been sampled or were the new mutations since the time of introgression. Analysis of non-silent mutations (**supplementary Fig. S13**) did not reveal a single causative mutation, but variant chr10: 29,439,011 has the highest correlation with the phenotype.

### Ear morphology influenced by introgressed *MSRB3*

Another prominent introgressed region with high *F*_ST_ contains MSRB3, encoding methionine sulfoxide reductase B3 (**Fig. 5A-C, supplementary Fig. S14**). Interestingly, ear morphology has been mapped to *MSRB3* in sheep using breeds fixed for divergent ear types (Paris et al., 2020), designated as ear size (large-eared vs. small-eared) and ear erectness (drop-eared vs. prick-eared). This gene yielded significant *f*_*d*_ values in 9 pairwise comparisons of argali vs. domestic population, encompassing chr3:154,000,001-154,090,000 (**Fig. 5A**). This was confirmed by the absolute divergence *d*_XY_ of argali and Oula, and of argali and Prairie Tibetan populations. (**Fig. 5B**), which indicated introgression rather than shared ancestry (ILS) (Martin, et al. 2015). By contrast, the *d*_XY_ of Iranian mouflon and either Oula or Prairie Tibetan populations are elevated (**supplementary Fig. S15**), indicating that the phylogenetic relationship of this region deviates from the phylogeny of the *Ovis* species.

**Fig. 5.**
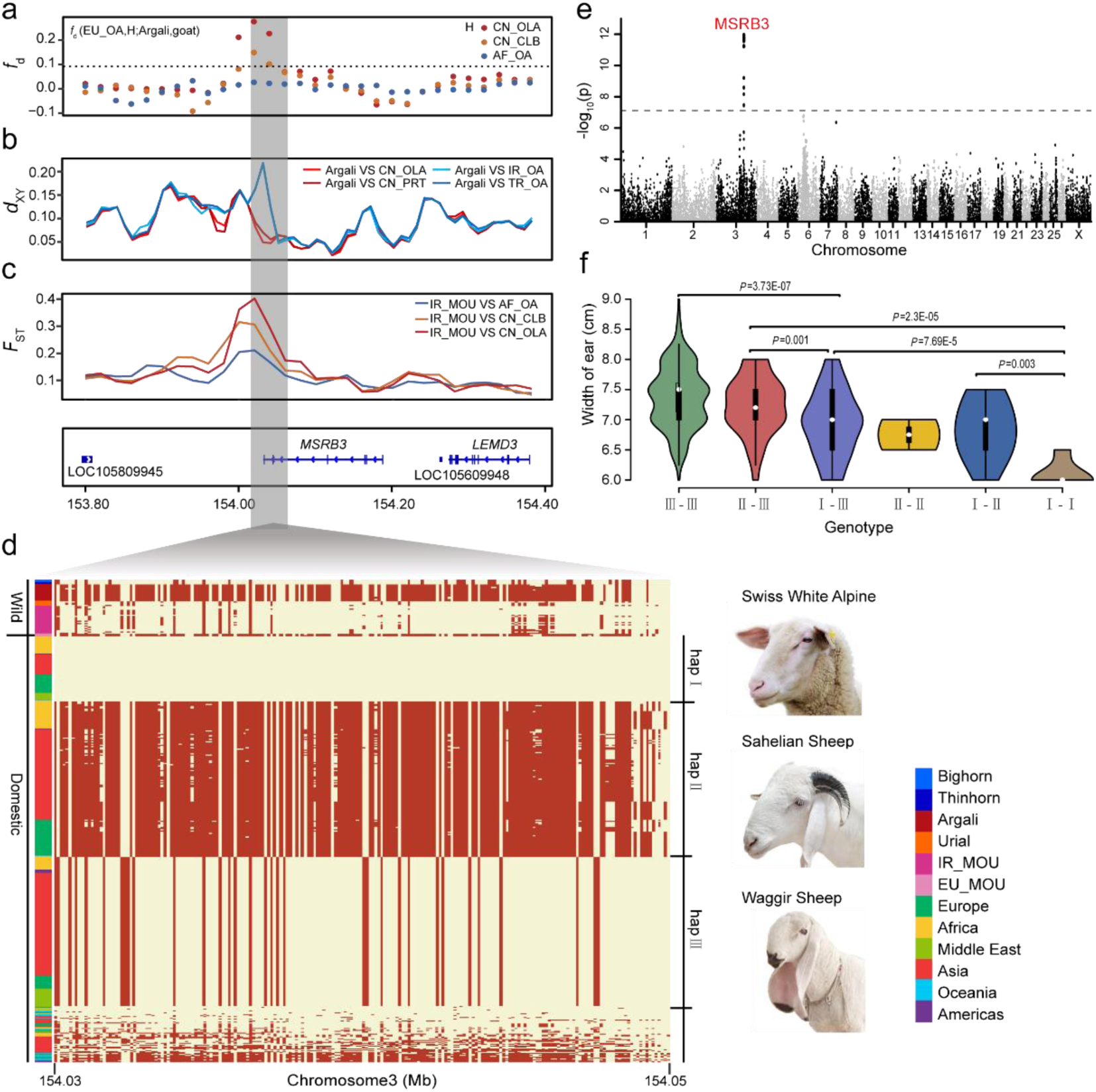
Identification and functional annotation of introgression segment at the *MSRB3* locus from argali to domestic sheep. (A-C) Distributions of *f*_*d*_ (EU_OA, H; Argali, Goat), *d*_XY_ values and *F*_ST_ surrounding the introgressed region (Oar_v4.0 chr3: 153,800,001-154,380,001). The horizontal dashed line in *f*_*d*_ track indicate the significance cutoff (*P*<0.05). (D)Haplotype pattern in the potential introgressed region (chr3: 154,030,048-154,062,195) of *MSRB3* gene. Each row is a phased haplotype, and each column is a polymorphic SNP variant. The reference and alternative allele are indicated in light yellow and red, respectively. Hap-II was the introgressed haplotype from argali. (E)GWAS -log10 *P* values for the width of the external ear of East-Friesian × Hu F2 crossbreds are plotted against position on the chromosomes. The gray horizontal dashed lines indicate the genome-wide significance threshold of the GWAS (7.72235E-08). (F)The violin plots refer to width of external ear for East-Friesian × Hu hybrids with different genotypes. III-III refers to homozygote of hap III defined in Fig. 5D, the other five genotypes are denoted accordingly.

A haplotype plot of the ∼32-kb *MSRB3* region across 1,167 individuals (**Fig. 5D, supplementary Fig. S16**) group into three main haplogroups, denoted as hap-I, hap-II and hap-III. These three haplogroups were corroborated by haplotype network and ML tree, in which domestic sheep haplotypes assigned to three clusters (**supplementary Fig. S18** and **S19**). The hap-II cluster is close to argali haplotypes (**supplementary Fig. S18** and **S19**), in consistent with it being introgressed from argali. Furthermore, hap-II has the highest frequency in domestic sheep (1109/1831, 0.605), and is fixed in Finnsheep (n=24), Hanzhong (n=10), Tibetan Oula (n=28), Feral (n=6) and Old Spael sheep, and nearly fixed (≥0.95) in Cameroon, Gotland, Ouessant sheep (**supplementary Fig. S17** and **supplementary Table S4**). Due to its high frequency among sheep breeds, we speculated this haplogroup was likely to confer an adaptive advantage over the other two groups. Intriguingly, all the European mouflon (n=3) in this study were likewise fixed for hap-II, suggesting that this introgression probably occurred before the first wave of migration of sheep into European(Chessa, et al. 2009; Lv, et al. 2015). The frequency of hap-I is relatively low across all domestic sheep (155/1831, 0.085), but has a high frequency in Swiss White Alpine (8/8,1), Mossi sheep (4/6, 0.67) and Diqing sheep (14/18, 0.78), all of which generally have small ears. Hap-III is found in breeds with exceptionally large and floppy ears (567/1831, 0.309), including Waggir (18/18, 1), Karakul (6/6, 1) and Duolang (63/68, 0.926), but also in Solognote (3/16, 0.188), Shetland (3/14, 0.214), Norwegian White (3/6, 0.5), Drente Heath (4/8, 0.5), East Friesian (15/20, 0.75) and Texel (6/6, 1) sheep that have small ears, suggesting that in addition to *MSRB3* other genes are involved in ear morphology.

For a more controlled analysis of a link between *MSRB3* variants and ear size, we used an F2 East-Friesian × Hu sheep hybrid population. We performed genome-wide association study (GWAS) of all the external ear traits, including measured width and length, in F2 hybrids (n=323) (**Fig. 5E, supplementary Fig. S20**). The ear width revealed a single significant association peak located in *MSRB3* (**Fig. 5E, supplementary Table S6**), but there was no significant signal associated with ear length (**supplementary Fig. S20**) or the other ear traits. Crossbred individuals with different haplotype combinations (**Fig. 3F**) or different genotypes of diagnostic SNPs (**supplementary Fig. S21, S22**) displayed significant difference in ear width.

### Complex patterns of introgressed regions within *VPS13B*

Another strong introgression signal was found in *VPS13B* (vacuolar protein sorting 13 homolog B), which showed the most significant introgressed signals from urial according to LAI (**Fig. 3B, supplementary Fig. S23**), as well as several consecutive outlier windows in the top 1% *f*_*d*_ values (**Fig. 3D** and **Fig. 6A**). *VPS13B* is a large gene spanning about 800 kb and has a complex structure with 50 exons and 6 alternatively spliced transcripts. It encodes a large protein with more than 4000 amino acids. The LAI results showed that there were two urial introgressed regions located in *VPS13B* chr9:77,117,407-77,437,296 (319.8 kb) and chr9:77,511,156-77,666,735 (155.5 kb) (**Fig. 3B**), comprising 4 and 3 major haplogroups respectively and covering about 59% of the gene (**Fig. 6E-F, supplementary Fig. S25, S26**). The haplogroups in these three regions form five major haplotype combinations, at least one of which is a recombinant (**Fig. 6D-F**). In addition, there is another separate introgression signal derived from mouflon in this gene chr9:76,946,737-77016,847, with three haplogroups (**Fig. 3B** and **6D, supplementary Fig. S24**), one of which is tightly linked to one of the five more downstream haplogroup combinations. The compound introgressed haplotype appears to have high frequency in Tibetan sheep (Oula: 0.9, Prairie Tibetan: 1; Valley Tibetan: 0.75), while at low frequency in domestic sheep from Iran (0.125), Turkey (0.045), America (0.063) and Australia (0).

**Fig. 6.**
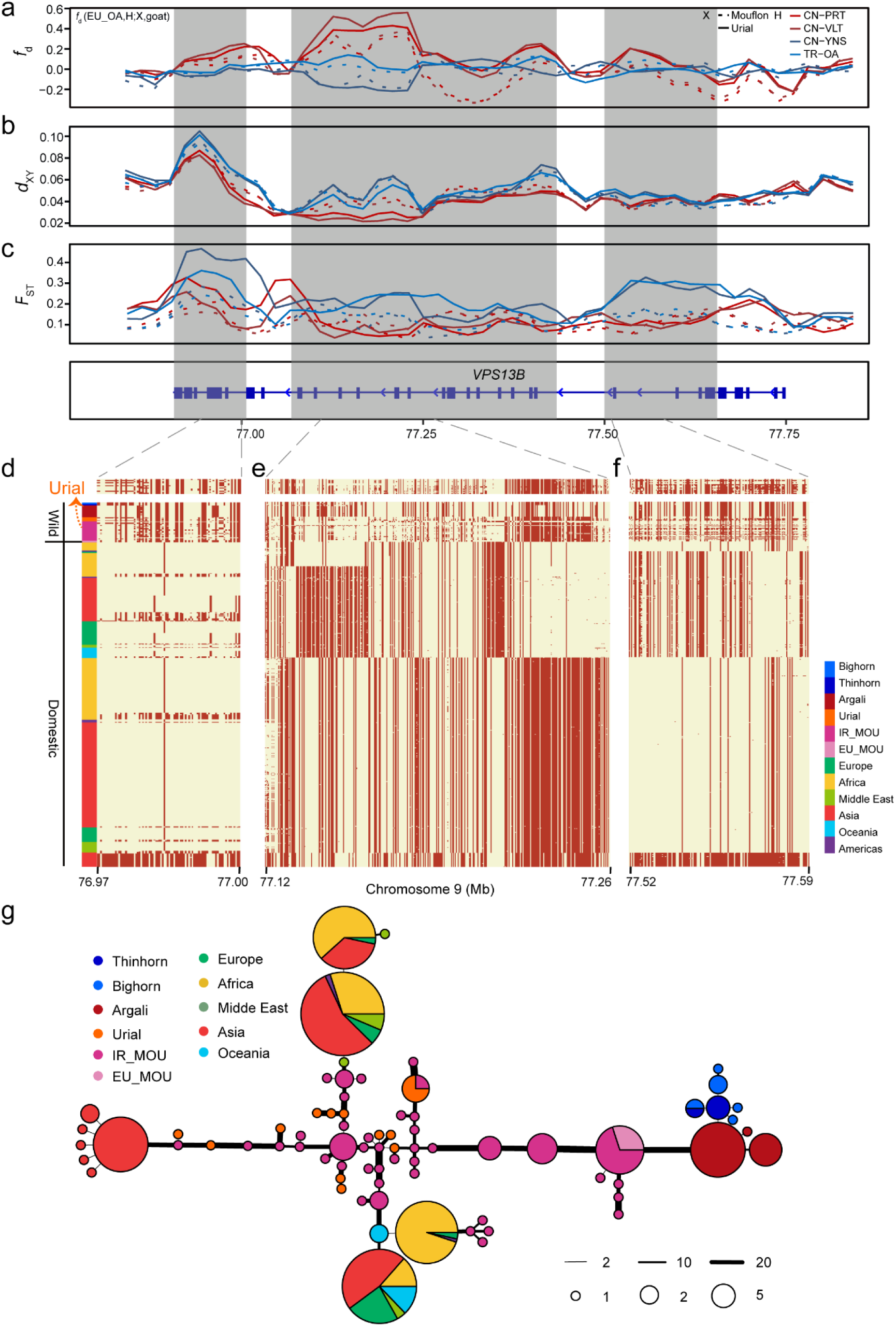
Signals of introgression in the *VPS13B* gene. (A-C) Distributions of *f*_*d*_ (EU_OA, X; Urial, Goat), *d*_XY_, and *F*_ST_ extending the three introgressed regions. The gene structures of *VPS13B* are indicated at the bottom track. (D-F) The patterns of haplotype sharing for part of introgressed regions. Each row is a phased haplotype, and each column is a polymorphic SNP variant. The reference and alternative alleles are indicated in light yellow and red, respectively. (G)A haplotype network (chr9:77145349 -77214813)generated by the R software package PEGAS based on 192 SNPs of 744 haplotypes.

These signals were also supported by *d*_XY_ and *F*_ST_ values which were lower between introgressed haplotypes and urial than mouflon, despite the closer phylogenetic position of the later to domestic sheep (**Fig. 6B-C, supplementary Fig. S27**). Whereas, this pattern was almost undetectable in partial introgressed region. We built haplotype networks of each region to investigate in detail the donor of introgressed haplotypes, but due to intermixed haplotypes we cannot distinguish whether the donor was urial or Iranian mouflon (**Fig. 6G, supplementary Fig. S28-30**). Similar to *RXFP2*, urial and Asiatic mouflon probably share the *VPS13B* haplotypes, which precludes an identification of the origin of the introgressions into domestic sheep.

*VPS13B* is functionally relevant to numerous phenotypes and diseases. It causes Cohen syndrome in humans with diverse manifestations including microcephaly, craniofacial and limb anomalies. In addition, variations in *VPS13B* affect face morphology, particularly nose morphology in human and mice(Bonfante, et al. 2021). Although the role of the introgressed fragments in *VPS13B* in sheep cannot at this stage be functionally verified, observations in other species suggest it may play a role in the development of facial shape.

## Discussion

In the present study, we performed a detailed investigation of introgression in sheep and evaluated the amounts of sequence introgressed from each wild relative into domestic sheep. We have focused on the most consequential breed-specific variants by selection of fragments of >100 kb with an *F*_ST_ > 0.1. We also present an in-depth investigation of three regions containing the genes *RXFP2, MSRB3* and *VPS13B*, which have been introgressed from wild sheep and now occur in a substantial proportion of the global sheep population. We show how these haplotypes are associated with variation in horn phenotypes and facial morphological variation in domestic sheep.

### Wild-domestic introgressions

In domestic sheep, we detected that the average proportions of wild relative sequence decrease with the phylogenetic distance of wild sheep species and domestic sheep (**Fig. 2A**). The strong signal of early sympatric gene flow of the Iranian mouflon into ancestral domestic sheep is geographically plausible and explains the high proportion of mouflon-derived sequences in domestic sheep. Domestic sheep have acquired urial DNA segments either directly from sheep breeds in the eastern distribution range of the urial (**Fig. 1**) or indirectly via the Iranian mouflon population(Demirci, et al. 2013). Subsequent dispersal has brought domestic sheep into contact with argali. The introgression from snow sheep had also been proved previously (Chen, Xu, et al. 2021b). A considerable genetic overlap of Asiatic mouflon and urial (Deng, et al. 2020; Chen, Xu, et al. 2021b) indicated incomplete speciation and/or mutual introgression. This has resulted in an incomplete differentiation of these species and does not allow a clear differentiation of Asiatic mouflon and urial as source of introgression of *RXFP2* and *VPS13B*.

Accurate identification of donor species may depend on the availability of whole genome sequencing (WGS) data from wild species candidates, and on method used to infer it. Hu et al. proposed argali introgression into *RXFP2*, but did not test the Asiatic mouflon(Hu, et al. 2019). Our data, especially the haplotype network (**Fig. 4C**), clearly indicate that, although the argali haplotype does resemble the introgressed haplotype (hap-c), hap-c has much closer affinity with haplotypes in Iranian mouflon. Moreover, hap-c is actually geographically widespread among domestic sheep, being found in both Tibetan, European and African sheep breeds, which has not been shown before (**Fig. 4D, supplementary Fig. S10** and **supplementary Table S4**). This supports that introgression of this haplotype predated the global dispersal from the sheep domestication rather than much later and localized to the Tibetan Plateau.

### Long divergent haplotypes contribute to diversity of sheep

We found that introgressed wild haplotypes covered about 8% of the sheep genome, and therefore contributed substantially to the diversity of domestic sheep, on the level of either individual or breed-specific variation. As indicated by **Fig. 3B**, we focused on a small proportion of all introgressed regions, but fragments that are shorter than 100 kb have a more random distribution across the breeds (low SD of within-breed frequencies) and do not appear to have been selected after domestication (low *F*_ST_ between all 16 breed groups and Asiatic mouflon). Despite this, they may still contribute to the overall diversity of domestic sheep.

Breed-specific introgression may well be related to local adaptation through their link to sheep phenotypes, e.g. hypoxia responses and high-attitude adaptation (Yang, et al. 2016; Hu, et al. 2019; Lv, et al. 2021), resistance to pneumonia (Cao, et al. 2020) and reproduction (Liu, et al. 2016). It would be a reasonable expectation that traits resulting from human selection (Xu and Li 2017; Li, Yang, Shen, et al. 2020) were only indirectly influenced by wild introgression, such as the different wool (Jackson, et al. 2020) and tail types (Kalds, et al. 2021). However, the absence of horns, a typical domestic feature, corresponding to *RXFP2* haplotypes is also detected in “horned” Asiatic mouflon. A testable hypothesis is that *RXFP2* of wild sheep is involved in balanced selection controlling the size of the horns.

A common feature of this study and comparable studies of cattle (Mei, et al. 2018; Wu, et al. 2018; Chen, Shen, et al. 2021a) and goats (Zheng, et al. 2020) are the observation of long (50 kb or longer) divergent haplotypes, associated with introgression or divergent selection (Via 2012). Similar divergence outliers, or genomic islands of differentiation have been observed for the human X chromosome (Shimada, et al. 2007); Neanderthal introgressions in the human genome (McArthur, et al. 2021); rattlesnakes (Dowell, et al. 2018); *C. elegans* (Lee, et al. 2021) and sunflowers (Todesco, et al. 2020). Divergence of homologous sequences inhibits recombination(Metzenberg, et al. 1991; Opperman, et al. 2004) which explains the absence of intermediates of the diverged haplotypes and allows to retain the divergence of the haplotypes.

In conclusion, using whole-genome sequencing data of large-scale individuals, we clarified the phylogenetic relationship among the eight extant species in the *Ovis* genus. In addition, we generated a global admixture graph of wild relative in diverse domestic sheep populations and determined whether positive selection had acted on these fragments. We also highlighted three introgressive regions in *RXFP2, MSRB3* and *VPS13B*. Through detailed haplotype and functional analyses, we evaluated the role of long divergent haplotypes from wild relatives in shaping the morphological traits of domestic sheep, which may be a ubiquitous phenomenon in animal evolution.

## Materials and Methods

### Sample collection

We newly sequenced 156 samples of WGS comprising 147 domestic sheep (*O. aries*) and 9 wild relatives (7 argali [*O. ammon*], 1 urial [*O. vignei*] and 1 European mouflon [*O. musimon*]) (**Fig. 1** and **supplementary Table S1** and **S2**). Following standard library preparation protocols, we used at least 0.5 μg of genomic DNA for each sample to construct paired-end library with insert sizes from 300 to 500 bp. Sequencing was performed on the Illumina HiSeq X Ten platform with a mean coverage of 13.30×. WGS data for 60 wild species (1 snow sheep [*O. nivicola*], 3 bighorn [*O. Canadensis*], 2 thinhorn [*Ovis dalli*], 13 argali, 6 urial, 33 Asiatic mouflon [*Ovis orientalis*] and 2 European mouflon), and 951 domestic individuals were obtained from previous studies (Yang et al. 2016; Alberto et al. 2018; Naval-Sanchez et al. 2018; Pan et al. 2018; Wang et al. 2018; Hu et al. 2019; Deng et al. 2020; Li et al. 2020; Upadhyay et al. 2020) (NCBI https://www.ncbi.nlm.nih.gov/, Nextgen http://projects.ensembl.org/nextgen/). The domestic samples originated from 154 different breeds with a geographic origin from Asia to the Middle East, Europe, Africa, Oceania and America (**supplementary Table S1**).

### Read alignment and variant calling

We firstly removed low-quality sequence reads of combined dataset by TRIMOMMATIC v.0.39 (BolgerLohse and Usadel 2014). Next, we aligned cleaned reads to Oar v.4.0 (https://www.ncbi.nlm.nih.gov/assembly/GCF_000298735.2) using the program BURROWS-WHEELER ALIGNER v.0.7.17 (BWA-MEM) algorithm (Li and Durbin 2010) with the default parameters. Duplicate reads were excluded using PICARD Markduplicates and bam files were sorted using PICARD SORTSAM (Picard v2.18.2 http://broadinstitute.github.io/picard/). Then, Genome Analysis Toolkit (GATK version 4.2.0.0) (McKenna et al. 2010) was performed to realign the reads around indels with REALIGNERTARGETCREATOR and INDELREALIGNER modules. To obtain the candidate SNPs from bam files, we used the workflow adapted from GATK HAPLOTYPECALLER to create genomic variant call format (gVCF) file for each sample. After merging all gVCF files, we implemented following criteria to SNPs using GATK VariantFiltration to avoid false-positive calls “ QD <2.0 || FS > 60.0 || MQRankSum <-12.5 || ReadPosRankSum < -8.0 || SOR >3.0 || MQ <40.0 “. SNPs not meeting the following criterias were further excluded: (1) biallelic variation; (2) missing rate < 0.1; (3) mean reads depth (DP) > 1/3× and < 3×. For remaining SNPs, imputation and phasing were simultaneous performed using BEAGLE v4.1 (Browning and Browning 2007; Browning and Browning 2016) with default parameters. SNPs and indels were annotated using the software ANNOVAR (WangLi and Hakonarson 2010).

### Population structure and phylogenetic analysis

We analyzed the population structure using 293 representative samples (Table S2), including all wild species and 56 domestic breeds. We used genome-wide 332,990 fourfold degenerate (4DV) sites to construct a maximum likelihood (ML) phylogenetic tree using RAxML v8.2.9 (Stamatakis 2014) with the following parameters: -f a -x 123 -p 23 -# 100 -k -m GTRGAMMA. The robustness of specific tree topology was tested by 100 bootstraps. The final tree topology was visualized using INTERACTIVE TREE OF LIFE (iTOL, Letunic and Bork 2016), and rooted at the branch of goat (**Fig. 2A**; **supplementary Fig. S1**).

PCA of whole-genome SNPs using was performed with the SMARTPCA program in the package of EIGENSOFT v.6.0beta (PattersonPrice and Reich 2006). To clarify the relationship between wild populations and domestic sheep, four groups were used for the PCA: (1) 293 samples, with the first two principal components cumulatively explaining 20.91% of the total variance. (2) 267 individuals including 3 European mouflon, 31 Asiatic mouflon and 233 domestic sheep; (3) 233 domestic individuals sampling from eight different regions, including Africa, Americans, Australian, China, South Asia, European, Iran and Turkey; (4) 117 individuals from 11 Chinese breeds (**Fig. 2B-D, supplementary Fig. S3**).

We used ADMIXTURE v1.23 (Alexander et al. 2009) to infer K=2 to K= 9 clusters of related individuals. For each K, we ran ADMIXTURE 20 times and calculated the mean cross-validation (CV) error to determine the optimal group number, the minimum CV value among 20 repetitions of each K was taken as the final result (**Fig. 2F**; **supplementary Fig. S4**).

### Selective sweep analysis

To detect potential selective signals, we calculated *F*_ST_ in the pairwise comparisons between Iranian mouflon and each of the domestic populations. The 233 domestic sheep were divided into 16 groups according to breeds and region of origin: EU_OA (Europe), AM_OA (America), AF_OA (Africa), TR_OA (Turkey), IR_OA (Iran), CN_YNS (Yunnan), CN_WZM (Ujimqin), CN_TAN (Tan), CN_STH (Small-tailed Han), CN_PRT (Prairie Tibetan), CN_VLT (Valley Tibetan), CN_OLA (Tibetan Oula sheep), CN_HU (Hu sheep), CN_CLB (Cele Black sheep), CN_BYK (Bayinbuluke sheep) and AU_MRN (Australian Merino). *F*_ST_ was calculated in 50-kb sliding windows with 20-kb step size (**supplementary Fig. S9, S14** and **S23**) using vcftools v.0.1.13 (Danecek et al. 2011). In each comparison, the top 1% genomic regions with the highest scores overlapped were considered to be potential selective signatures.

### Whole-Genome Analysis of Genomic Introgression

#### Estimation of introgression on population scale

We implemented *D*-statistics with DSUITE (Malinsky et al. and Svardal 2021) across all combinations of the 16 species/populations defines as described above. The species/population tree required for DSUITE was constructed using Treemix (Pickrell and Pritchard 2012) without assuming gene flow (-m 0) using goat as outgroup (**supplementary Fig. S5**). Then, D and *f*4-ratios of all populations were calculated with the DTRIOS module, and the results for each chromosome were combined with Dtrioscombine module. After that, D and *f* statistics were calculated for each branch of the population tree using the FBRANCH module, and visualized the statistical results using dtools.py script provided in the DSUITE software (**supplementary Fig. S5**). Because there were few alleles sharing between domestic sheep from Europe and the other wild species and domestic populations (**supplementary Fig. S5**), domestic European samples were identified as non-introgressed reference population.

#### Identification and localization of genomic introgression

We used local ancestry inference (LAI) implemented in LOTER (Dias-Alves, et al. 2018), which uses phased data and has been shown to outperform other tools for more ancient admixture. We specified seven wild relatives (1 snow sheep, 3 bighorn and 2 thinhorn, 12 argali, 6 urial, 31 Iranian mouflon, 3 European mouflon), and European domestic sheep as reference population, in which European domestic sheep (n=30) was the control population for domestic component. It was assumed that a haplotype of an admixed domestic individual consists of a mosaic of existing haplotypes from the eight reference populations. For each fragment, LOTER derives the most likely ancestral origin on the basis of allele frequencies of reference populations and the selected populations. We calculated introgression percentages from each of the wild relatives into the haploid genomes (**Fig. 3A**) and merge overlapping introgressed regions from the same source. Then, the frequencies in the 16 domestic groups with their standard deviations (SD) and ranges (max-frequency minus min-frequency) were calculated for each selected fragment (**Fig. 3B**).

#### f_d_ in sliding windows

We computed the modified f-statistic (*f*_*d*_) value (Martin et al. 2015) using a 50-kb sliding window with 20-kb step size in the form of *f*_*d*_ (EU_OA, domestic populations; wild species, goat), where EU_OA represents the European domestic sheep (n=30) and domestic populations include 16 populations described above. We evaluated the statistical significance using two-tailed Z-test. We calculated the *P* values according to Z-transformed *f*_*d*_ values, and the windows with *P* < 0.05 was defined as potential introgressed regions (**Fig. 3C-E**). Mean pairwise sequence divergence (*d*_XY_) (Martin et al. 2015) was also calculated for 50-kb windows with 20-kb steps across whole genome using same populations above (**Fig. 5B, 6B**; **supplementary Fig. S15**).

#### Incomplete Lineage Sorting (ILS)

In order to exclude common ancestry as explanation for the presence of introgressed fragments, we calculated the expected length of ancestral sequence shared by domestic sheep and each wild relative, respectively. The expected shared ancestral sequence length (*L*) is calculated as *L*=1/(*r*×*t*), in which *r* is the recombination rate per generation per base pair (bp), and *t* is the length between wild relatives and domestic sheep since divergence. The probability of a length of at least m is 1-*GammaCDF* (m, shape = 2, rate =1/*L*), in which *GammaCDF* is the Gamma distribution function and the numbers within the parenthesis are its arguments (Huerta-Sánchez et al. 2014). We used a generation time of 4 years (Guerrini et al. 2015), a combination rate of 1.0×10^−8^ (Kijas et al. 2012) and the following divergence times: 0.032 Ma between Iranian mouflon and domestic sheep, 1.26 Ma for urial and domestic sheep, 2.36 Ma for argali and domestic sheep, and 3.12 Ma for bighorn (or thinhorn) and domestic sheep (Bunch et al. 2006; Rezaei et al. 2010; Yang et al. 2017; Li et al. 2021). This gives expected lengths of *L* (Iranian mouflon) = 6,192 bp, L (urial) = 159 bp, *L* (argali) = 85 bp, and *L* (bighorn/thinhorn) = 64 bp. We then removed inferred introgressed fragments shorter than *L*, and calculated the total length of remaining introgressed tracks. The length distributions are showed in **supplementary Fig. S6, S7**. Probabilities of length of observed introgressed regions calculated by the R function pgamma are 0.00493 for 46.10 kb (*RXFP2* introgressed region) and zero for 31.70 kb (*MSRB3* argali-introgressed region), 319.89 and 155.58 (*VPS13B* urial-introgressed regions), and 0.000149 for 70.11 kb (*VPS13B* mouflon-inrogressed).

### Haplotype patterns and network

To view the specific genotypes patterns of the prominent introgressed regions, including *RXFP2, MSRB3* and *VPS13B*, we extracted the phased SNPs in these regions from 1,167 whole-genome sequencing individuals and visualized specific genotypes patterns in a heatmap (**supplementary Fig. S10, S16, S24-26**). We also constructed haplotype networks of *RXFP2, MSRB3 and VPX13B* using R package PEGAS (Paradis 2010) based on the pairwise differences (**Fig. 4D, 6G**; **supplementary Fig. S19, S27-29**). We screened and eliminated samples whose haplotypes were interrupted due to recombination and removed SNPs with minor allele frequency ⩽5%. In the 46.3 kb *RXFP2* introgressed region we retained 333 samples from 6 wild species and 20 domestic sheep breeds and 221 SNPs. In the 23 kb *MSRB3* introgressed region we retained 202 SNPs in 201 individuals. We analyzed the sheep ear shapes of the corresponding varieties of three haplotypes, and defined three main haplotypes as hap-I, hap-II and hap-III. We constructed an ML tree based on the introgressed region located in *RXFP2* and *MSRB3* respectively (Fig. S12 and S18).

### Genome-wide association study

From an East-Friesian sheep × Hu Sheep F2 generation bred by Gansu Yuansheng Agriculture and Animal Husbandry Technology Co.,Ltd. (Jinchang, Gansu) 323 samples were collected. Phenotypes include ear length and width, birth weight and age. DNA was collected from blood samples and whole genomes were sequenced by Shijiazhuang Boruidi Biotechnology Co., Ltd (Shijiazhuang, Hebei) using 40K liquid chip generated by genotyping by target sequencing. Raw fastq files were filtered using fastp (Chen et al. 2018b), reads were mapped to Oar_rambouillet_v1.0 and the variation were summarized in a VCF file. We used PLINK1.9 (Purcell et al. 2007) to remove samples with > 10% missing genotypes and SNPs with minor allele frequency < 0.05 and >10% missing scores, retaining 317 sheep and 209,625 SNPs. To improve variant density, we used BEAGLE5.0 (Browning et al. 2018) to impute genotype using reference panel size of 43 East Friesen sheep and 8 Hu sheep with default settings and removed SNP with DR^2^ (dosage R-squared) ⩽ 0.8, resulting in a total of 647,471 SNPs.

GWAS was conducted using GEMMA(0.98.3) (Zhou and Stephens 2012) with the linear mixed model and the model:

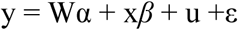

*y* is the phenotype of *n ×1* vector; *W* the *n×c* matrix of covariates including fixed effects; *α* the *c-*vector of the corresponding coefficients including the intercept; x is the *n-*vector of markers; *β* the effect size of the markers; u the *n-*vector of the random effect with u∼MVN_n_(0, *λτ*^-1^*K*); MVN_n_ the n-dimensional multivariate normal distribution; *λ* the ratio between the two variance components;*τ*^-1^ is the variance of the residual errors; *K* represents the known *n×n* relatedness matrix calculated by SNP markers; *ε* the random error *n*-vector with *ε*∼ MVN_n_ (0, *τ*^-1^*I*_*n*_), where *I*_*n*_ denotes *n*×*n* identity matrix. The genome-wide significance threshold was set to be 7.72235E-08 (0.05/647,471) after the Bonferroni correction.

## Acknowledgments

This work was supported by grants from the National Natural Science Foundation of China (U21A20247 and 31822052) Shaanxi Innovation Team Project (2022TD-10), Natural Science Basic Research Program of Shaanxi (2021JCW-11) to Y.J. and Research on High-efficiency and Healthy Breeding Technology for Both Milk and Meat Sheep, the Jinchang Meat Sheep Test Demonstration Base (TGZX202137) to Y.S. We thank the High-Performance Computing platform of Northwest A&F University for providing computing resources. We express our thanks to the owners of the sheep for providing samples (see **supplementary** Table S2).

## Author Contributions

Y.J. lead the project, and designed and conceived the study. H. C., J.W., and Z.Z. performed the data analysis. Z.Z., J.W., J.S. and M.B. collected GWAS sheep samples. M.L., W.L., S.H., Y.S. and L.Z. collected local sheep samples. M.L., Y.C., and Y.G. assisted in data interpretation. H.C. prepared the manuscript. Y. J., J.A.L., R. H., R. L., M.L. and X. W revised the manuscript.

### Declaration of interests

The authors declare no competing interests.

### Ethics statement

Blood samples were taken by conforming with the Helsinki Declaration of 1975 (as revised in 2008) concerning Animal Rights, and this study was reviewed and approved by the Animal Ethical and Welfare Committee (DK2021019), Northwest A&F University, China.

### Data Availability

Raw sequencing data generated in this study have been deposited to the NBCI BioProject database under accessions numbers PRJNA814428 and PRJNA521847 (Tibetan sheep).

### Supplemental information

Figures S1-S31

Table S1-S6

## Notes

### Competing Interest Statement

The authors have declared no competing interest.

